# Spi-C and PU.1 counterregulate *Rag1* and *Ig*κ transcription to effect immunoglobulin kappa recombination in small pre-B cells

**DOI:** 10.1101/2022.12.28.522117

**Authors:** Li S. Xu, Jiayu T. Zhu, Hannah L. Raczkowski, Carolina R. Batista, Rodney P. DeKoter

**Author notes:** Department of Microbiology & Immunology, Schulich School of Medicine & Dentistry, Western University, London, Ontario, Canada N6A 5C1.

## Abstract

B cell development requires the ordered rearrangement of *Ig* genes encoding H and L chain proteins that assemble into BCRs or Abs capable of recognizing specific Ags. *Ig*κ rearrangement is promoted by chromatin accessibility and by relative abundance of RAG1/2 proteins. Expression of the E26-transformation-specific (ETS) transcription factor Spi-C is activated in response to double-stranded DNA breaks (DSBs) in small pre-B cells to negatively regulate pre-BCR signaling and *Ig*κ rearrangement. However, it is not clear if Spi-C regulates *Ig*κ rearrangement through transcription or by controlling RAG expression. In this study, we investigated the mechanism of Spi-C negative regulation of *Ig*κ light chain rearrangement. Using an inducible expression system in a pre-B cell line, we found that Spi-C negatively regulated *Ig*κ rearrangement, *Ig*κ transcript levels, and *Rag1* transcript levels. We found that *Ig*κ and *Rag1* transcript levels were increased in small pre-B cells from *Spic*^-/-^ mice. In contrast, *Ig*κ and *Rag1* transcript levels were activated by PU.1 and were decreased in small pre-B cells from PU.1-deficient mice. Using chromatin immunoprecipitation analysis, we identified an interaction site for PU.1 and Spi-C located in the *Rag1* promoter region. These results suggest that Spi-C and PU.1 counterregulate *Ig*κ transcription and *Rag1* transcription to effect *Ig*κ recombination in small pre-B cells.

## Introduction

B cell development requires the ordered rearrangement of *Ig* genes encoding H and L chain proteins that assemble into BCRs or Abs capable of recognizing specific Ags. *Ig* rearrangement proceeds in a developmental sequence in which *IgH* V(*D*)J recombination precedes Igκ or Igλ V-J recombination. Igκ/λ V-J recombination occurs in small pre-B cells that have previously rearranged a functional *IgH* allele, have ceased proliferation in response to IL-7, and have turned on high expression of recombinase activating gene (RAG) proteins RAG1 and RAG2 (1, 2). *Ig*κ rearrangement is promoted by chromatin accessibility, regulated through transcription; and by relative abundance of RAG1/2 proteins (3).

*Ig*κ accessibility and transcription are controlled by two locus-specific enhancers, the intronic enhancer and the 3’ enhancer (4). *Ig*κ chromatin accessibility correlates with germline transcription initiating upstream of the Jκ1 segment – this transcript is alternatively named k^0^ or germline kappa-1 (*Glk1*) (5, 6). The 3’ enhancer contains a ETS transcription factor binding site that regulates B cell versus T cell specificity of Vκ-Jκ joining (7). *Ig*κ transcription and accessibility is repressed by STAT5 interaction with the *Ig*κ intronic enhancer, downstream of IL-7 receptor signaling in pro-B cells and large pre-B cells (8, 9).

Regulation of RAG protein abundance is an important second mechanism for regulation of *Ig*κ recombination. RAG1 and RAG2 together form the recombinase that catalyzes V(*D*)J recombination by recognition and cleavage of recombination signal sequences (10). Appropriate developmental regulation of *Rag1/Rag2* is essential to prevent off-target mutations leading to leukemia or lymphoma (11). RAG1/2 protein levels are regulated by specific degradation linked to the cell cycle (12). The closely linked *Rag1* and *Rag2* genes are also regulated transcriptionally (13). Several regulatory regions or enhancers of the *Rag* locus have been identified including Erag (14), R-Ten, and R1B/R2B (15). These regulatory regions interact with lineage-specific transcription factors to activate or repress *Rag* transcription at specific developmental stages (15). RAG-induced DNA double-stranded breaks (DSBs) promote allelic exclusion by signaling through ataxia telangiectasia mutated (ATM) kinase and the NF-κB essential modulator (Nemo) to repress *Rag* and *Ig*κ transcription (16).

One of the genes activated by DSBs in small pre-B cells through the ATM/NF-kB signaling pathway is *Spic* encoding the E26-transformation-specific transcription factor Spi-C. Spi-C is a lineage-instructive transcription factor that is important for the generation of myeloid and lymphoid cells (17). Induction of Spi-C in pre-B cells functions to repress cell cycle progression and *Ig*κ recombination by interacting with and repressing the *Syk, Blnk*, and *Ig*κ genes (18, 19) Spi-C is required for the development of splenic red pulp macrophages (20, 21). Spi-C is dynamically regulated in response to growth factor signals to regulate gene expression in B cells (22, 23).

Spi-C is highly related to the ETS-family transcription factors PU.1 and Spi-B that function as complementary transcriptional activators of BCR signaling and *Ig*κ rearrangement (24). Combined deletion of *Spi1* and *Spib* genes encoding PU.1 and Spi-B in mice results in a block to B cell development at the small pre-B cell stage followed by development of precursor B cell acute lymphoblastic leukemia (25, 26). PU.1 and Spi-B function as transcriptional activators while Spi-C functions as a transcriptional repressor by competing for PU.1 binding sites (27–29). In addition, DSBs induced by RAG activate Spi-C to displace PU.1 binding from the *Ig*κ 3’ enhancer to repress transcription (19). Therefore, PU.1/Spi-B and Spi-C may function to oppose one another’s function to regulate BCR signaling and *Ig*κ rearrangement during B cell development.

In this study, we investigated the hypothesis that Spi-C and PU.1 function as regulators of *Ig*κ recombination by regulation of *Rag* transcription as well as *Ig*κ transcription. To do this, we constructed a doxycycline-inducible Spi-C expression system in 38B9 pre-B cells. Induction of Spi-C inhibited *Ig*κ recombination and inhibited *Ig*κ and *Rag* mRNA transcription in 38B9 cells. To determine if Spi-C is required to repress *Ig*κ and *Rag* mRNA transcription during B cell development, we generated *Spic*^-/-^ mice by intercrossing *Spic*^+/-^ mice onto a mixed C57Bl/6 and 129S/v background. *Ig*κ and *Rag* mRNA transcripts were upregulated in small pre-B cells enriched from *Spic*^-/-^ mice relative to wild-type pre-B cells. Finally, we used chromatin immunoprecipitation to identify a regulatory binding site for PU.1 and Spi-B upstream of the *Rag1* gene. These results indicate that Spi-C and PU.1 are counterregulators of *Rag* and *Ig*κ transcription during B cell development.

## Materials and Methods

### Mice

*Spic*^+/-^ mice were maintained as previously described on a C57Bl/6 background (30). To generate *Spic*^-/-^ mice, *Spic*^+/-^ mice were mated to 129Sv mice (Charles River Laboratories, Pointe-Claire, Qc, Canada) to generate F1 C57Bl6/129Sv *Spic*^+/-^ mice. F1 mice were intercrossed to generate F2 *Spic*^-/-^ mice. All F2 *Spic*^-/-^ mice used for experiments used F2 *Spic*^+/+^ or *Spic*^+/-^ littermate mice as controls. All mice were housed in a specific pathogen free animal facility and monitored under an approved animal use protocol approved by the Western University Animal Care Committee.

### Cell Sorting

Bone marrow (BM) cells were prepared from euthanized mice and erythrocytes were removed using ammonium-chloride-potassium lysis. Erythrocyte-depleted BM cells were stained with antibodies including PE conjugated anti-CD19 (6D5, Biolegend, San Diego, CA), allophycocyanin conjugated anti-B220 (RA3-6B2, eBioscience, San Diego, CA), brilliant violet 421-conjugated anti-B220 (RA3-6B2, Biolegend), APC-conjugated anti-IgM (II/41, BD Biosciences, Franklin Lakes, NJ), FITC-conjugated anti-CD24 (M1/69, Biolegend), PE-conjugated anti-BP-1 (BP-1, BD Biosciences), biotin-conjugated anti-CD43 (S7, BD Biosciences), and PE/Cy7-conjugated streptavidin (Biolegend). Infected cultured cells were sorted based on green fluorescent protein (GFP) expression. Dead cells were distinguished using Sytox blue dead cell stain (Thermo-Fisher Scientific) or 7-AAD viability stain (Thermo-Fisher). Cell sorting was performed using a FACSAriaII instrument. Flow cytometry analysis was performed using a FACsCanto™ instrument (BD Immunocytometry systems, San Jose, CA). Flow data were analyzed using FlowJo 9.1 (Tree Star, Ashland, OR). The purity of sorted cells was determined to be ≥98%.

### Cell Culture

The 38B9 pre-B cell line (31) was cultured in Iscove’s Modified Dulbecco’s Medium (IMDM, Wisent, QC, Canada) supplemented with 10% FBS (Wisent), 1x penicillin/streptomycin/ L-glutamine (Wisent), and 5×10^-5^ M 2-mercaptoethanol (Sigma-Aldrich, St. Louis, MO). Cell lines were maintained in 5% CO2 atmosphere and 37°C. To induce differentiation and Igκ V-J rearrangement, 38B9 cells were treated with 0.5μM Imatinib (Sigma-Aldrich) for 48 hours (32). PU.1-inducible i660BM cells were cultured as previously described (33). Mouse bone marrow cells were cultured in complete IMDM medium supplemented with 5% supernatant from the J558L-IL7 cell line (34). For Interleukin-7 withdrawal experiments, cells were grown at 2×10^6^ /ml with IL-7 for 8 days, washed free of IL-7, and cultured without IL7 for 48 hours.

### Construction of a Spi-C Inducible System

Murine 3XFLAG-tagged Spi-C cDNA (35) was PCR amplified and ligated into the pRetro-Tre3G (Tet-regulatable element third generation) plasmid (Thermo-Fisher Scientific, Waltham MA). MIG-Tet3G was previously described (24). pRetro-Tre3G and MIG-Tet3G retrovirus was produced by transient transfection of Plat-E retroviral packaging cells (36). 38B9 cells were infected by “spinoculation” followed by enrichment for green fluorescent cells using cell sorting. Sorted cells were grown continuously in media supplied with 1μg/ml puromycin (Biobasic, Markham, ON) for two weeks for selection of resistant cells. The resulting cell line was named “inducible 38B9” (i38B9). To induce SpiC overexpression, i38B9 were treated with Doxycycline (Biobasic) at 1.5 μg/ml.

### Polymerase Chain Reaction

RNA was prepared from cultured or sorted cells using the RNeasy kit (Qiagen, Germantown MD) and cDNA was synthesized using the iScript cDNA synthesis kit (Bio-Rad, Hercules CA) following the manufacturer’s instructions. Reverse transcriptase quantitative polymerase chain reaction (RT-qPCR) was performed using SensiMix SYBR No-Rox master mix (Froggabio, Toronto, Canada) on a QuantStudio5 instrument (Applied biosystems, Thermo Fisher Scientific). Relative frequencies of mRNA transcripts were all determined as ratios compared to mRNA levels of beta-actin. Genomic DNA was extracted using the Wizard genomic DNA purification kit (Promega, Madison, WI). 500ng DNA from each sample was used for PCR. For amplification of Igκ V-J rearrangements from cDNA (37), 50 μl PCR reactions included magnesium chloride at 1.5mM, V_k_ primer at 3.2 μM, J_k_ primer at 0.2 μM, and 0.5 units of Takara LA-Taq™ DNA polymerase (Takara Bio Inc., Kusatsu, Shiga, Japan). PCR cycle was 95°C for 4 minutes, followed by 31 cycles at 95°C denaturing for 1 minute, 62°C annealing for 1 minute, and 72°C extension for 1 minute and 45 seconds, and a final 10-minute extension at 72°C. Primer sequences are described in **Supplemental Table 1.**

### Chromatin Immunoprecipitation

Spi-C was induced for 48hr in i38B9 cells with 1.5 μg/ml doxycycline, followed by crosslinking using 1% formaldehyde (Millipore-Sigma) and halted using glycine. Crosslinked pellets of up to 1 x 10^7^ cells were flash-frozen in liquid nitrogen prior to sonication. Frozen fixed pellets were resuspended in lysis buffer supplemented with HALT protease inhibitor (Thermo-Fisher) and sonicated for 25 cycles using the Bioruptor UCD-300 (Diagenode, Sparta, NJ). Immunoprecipitation of 3XFLAG-Spi-C bound chromatin was performed using anti-FLAG M2 magnetic beads (Millipore-Sigma). DNA was eluted from input chromatin or immunoprecipitated chromatin by heating. Eluted DNA was purified using the ChIP DNA clean and concentrator kit (Zymo Research, Irvine CA). qPCR analysis was performed using primers described in **Supplemental Table 1**.

### Statistical analysis

Statistical analysis was performed using Prism 9.4.1 (Graphpad Software, La Jolla, CA). Data presented as Mean ± SEM unless otherwise indicated. Statistical significance was determined using One Sample Wilcoxon test or Student’s t-test. All data points in figures in this study represent independent biological replicate experiments.

## Results

### Induced expression of Spi-C inhibits Ig kappa rearrangement in an inducible cell line

To study the role of Spi-C in *Ig*κ transcription and rearrangement, we undertook to construct a Spi-C inducible system using the Abelson (Abl) kinase-transformed pre-B cell line 38B9 (31). Cell cycle arrest, *Ig*κ V-J rearrangement, and B cell differentiation can be induced in 38B9 cells using the Abl kinase inhibitor imatinib (also known as STI571 or Gleevec) (32). We constructed a two-vector doxycycline-inducible system for ectopic expression of 3XFLAG-tagged murine Spi-C in 38B9 cells (Fig. 1A). With this system, Spi-C expression could be induced in inducible 38B9 (i38B9) cells up to 10-fold at 24 or 48 hours (Fig. 1B). Induction of Spi-C did not affect cell cycle progression or induce *Ig*κ recombination in i38B9 cells (data not shown).

**FIGURE 1.**
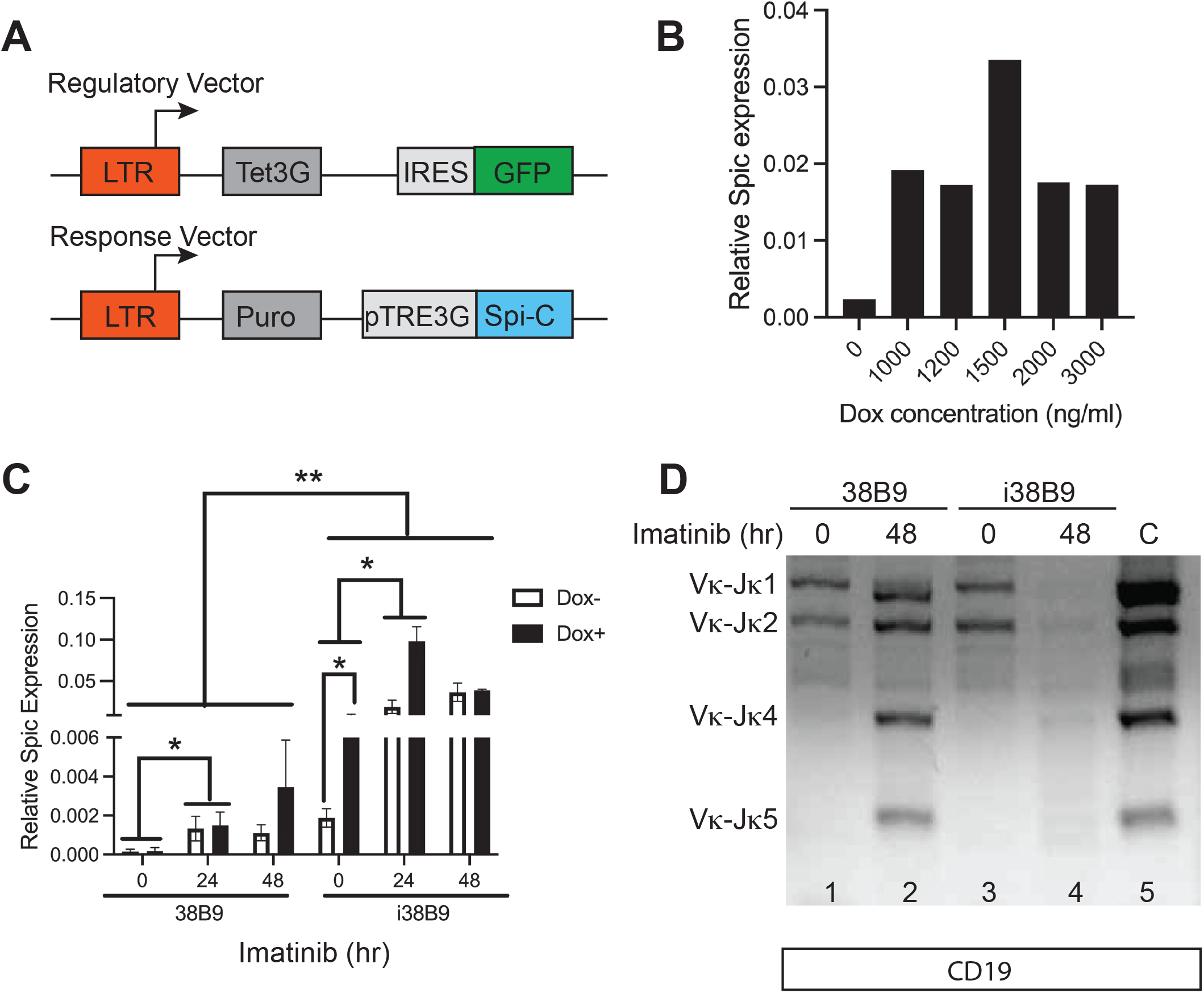
*Ig*κ rearrangement is inhibited by Spi-C in an inducible cell line. (**A**) Schematic of two-vector inducible system. Top panel shows regulatory vector, bottom panel shows response vector. LTR, long terminal repeat; Tet3G, third-generation tetracycline-regulatable activator; IRES, internal ribosomal entry site; GFP, green fluorescent protein; Puro, puromycin resistance gene; pTRE3G, third-generation tetracycline regulatable element. (**B**) Induction of *Spic* expression by doxycycline in inducible 38B9 (i38B9) cells. Doxycycline (Dox) concentrations in ng/ml are indicated on the x-axis, Relative *Spic* expression (*Spic*/β-*actin* ratios) are indicated on the y-axis. (**C**) Induction of *Spic* expression by doxycycline and imatinib in i38B9 cells. Times after treatment with 0.5 μM imatinib are shown on the x-axis. Induction with 1.5 μg/ml doxycycline is indicated by open or filled bars. Relative *Spic* expression (*Spic*/β-*actin* ratios) is indicated on the y-axis. (**D**) Inhibition of *Ig*κ V-J rearrangement 48 hr after imatinib treatment and Spi-C induction. PCR analysis was performed on genomic DNA prepared from cells treated with 0.5 μM imatinib at times indicated on the top as well as 1.5 μg/ml doxycycline. Positive control (WT) is from genomic DNA prepared from wild-type splenic B cells. \**p*<0.05 and ** *p*<0.01 using two-way ANOVA.

Next, 38B9 cells or i38B9 cells were treated with imatinib to induce *Ig*κ V-J rearrangement. Imatinib treatment induced 38B9 cell cycle arrest by 48 hours, as shown by flow cytometric analysis of a shift from high forward scatter to low forward light scatter suggesting a reduction in cell size (Supplemental Fig. 1A). *Spic* was expressed at low levels in untreated 38B9 or i38B9 cells and was increased substantially by 0.5 μM imatinib treatment (Fig. 1C). Prior to imatinib treatment, both 38B9 and i38B9 cells had detectable V_k_-J_k_1 and V_k_-J_k_2 rearrangements but undetectable Vκ-Jκ4 and Vκ-Jκ5 rearrangements (Fig. 1D). Imatinib treatment induced Vκ-Jκ rearrangements in both 38B9 and i38B9 cells (Supplemental Fig. 1B).

Next, the combined effects of inducing Spi-C using doxycycline and inducing *Ig*κ rearrangement with imatinib were investigated. Interestingly, imatinib increased levels of *Spic* mRNA transcripts even in doxycycline untreated 38B9 or i38B9 cells, suggesting activation of the *Spic* gene (Fig. 1C). However, *Spic* mRNA levels were induced over 50-fold by combined doxycycline and imatinib treatment (Fig. 1C, right side). Induction of Spi-C in i38B9 cells strongly inhibited imatinib-induced Vκ-Jκ rearrangements at the 48-hour timepoint (Fig. 1D, right panels). This result suggests that ectopic expression of Spi-C negatively regulates *Ig*κ rearrangement in pre-B cells.

### Repression of Ig kappa germline and Rag1 transcripts by Spi-C

*Ig*κ rearrangement might be inhibited by Spi-C through repression of *Ig*κ transcription, repression of RAG1/2 expression, or a combination of both. To determine the mechanism of inhibition of *Ig*κ rearrangement by Spi-C, gene expression was examined using RT-qPCR. *Rag1* and *Rag2* encode the recombinase-activating 1 and 2 proteins, while *Glk1* encodes an *Ig*κ germline “sterile” transcript that correlates with locus accessibility (6). *Glk1* but not *Rag1* or *Rag2* transcripts were inhibited by Spi-C at the 24-hour time point (Fig. 2A-C). However, *Rag1, Rag2*, and *Glk1* mRNA transcript levels were all reduced at 48 hours post-imatinib treatment in doxycycline-induced i38B9 cells (Fig. 2A-C). Therefore, ectopic expression of Spi-C inhibits *Ig*κ germline transcription, *Rag1*, and *Rag2* transcription in 38B9 cells.

**FIGURE 2.**
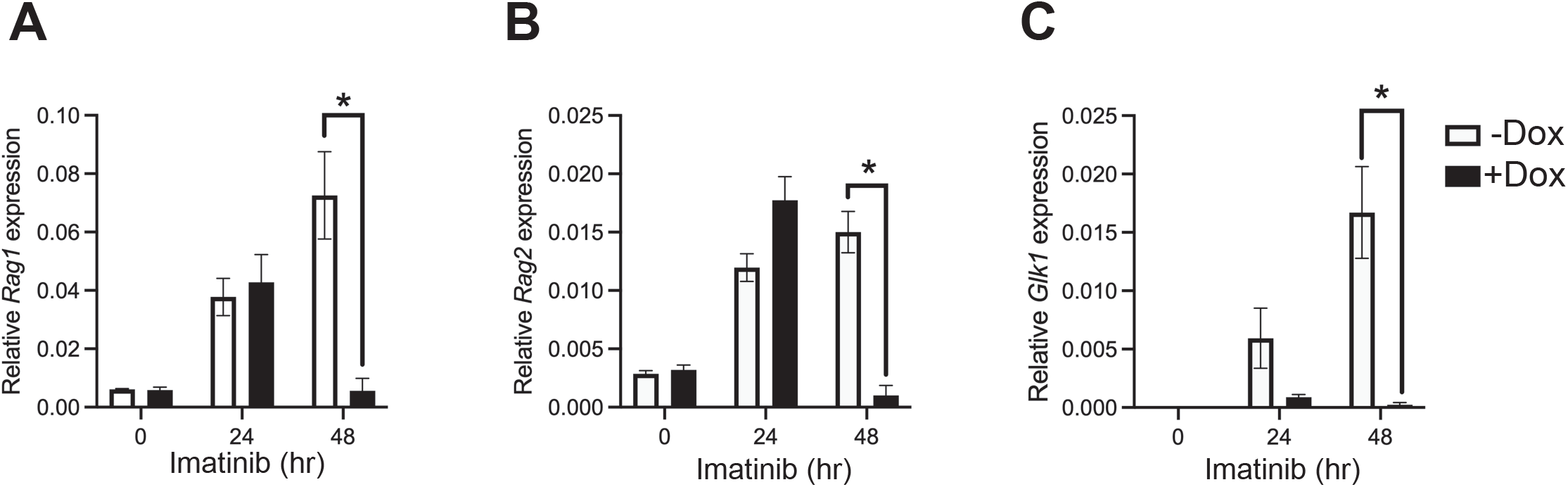
Repression of *Ig* kappa germline and *Rag1* transcripts by Spi-C. RNA was prepared from i38B9 cells treated for 0, 24, or 48 hr with 0.5 μM imatinib and induced or not with 1.5 mg/ml doxycycline (open versus filled bars). RT-qPCR was performed for the genes indicated on the y-axis. Relative expression (gene/β-*actin* ratios) is indicated on the y-axis. \**p*<0.05 using one-sample t and Wilcoxon test.

### Early B cell development is not altered in Spic^-/-^ mice

We set out to determine the effect of *Spic* loss-of-function on gene expression in developing pre-B cells in mice. *Spic*^-/-^ mice are born at low frequency due to preimplantation lethality of undetermined cause (20, 38). We found that *Spic*^-/-^ mice on a C57Bl/6 strain background were rarely generated, either at 3 weeks of age or fetal 14.5 days post-implantation (Table 1). However, by intercrossing F1 progeny of C57Bl/6 *Spic*^+/-^ x 129Sv mice, we were able to obtain *Spic*^-/-^ mice with a frequency of 14.8% by 3 weeks of age (Table 1, Fig. 3A). This outcome suggests that *Spic*^-/-^ mice on a mixed C57Bl/6 and 129.Sv strain background have reduced preimplantation lethality. This procedure allowed generation of *Spic*^-/-^ mice for further analysis.

**FIGURE 3.**
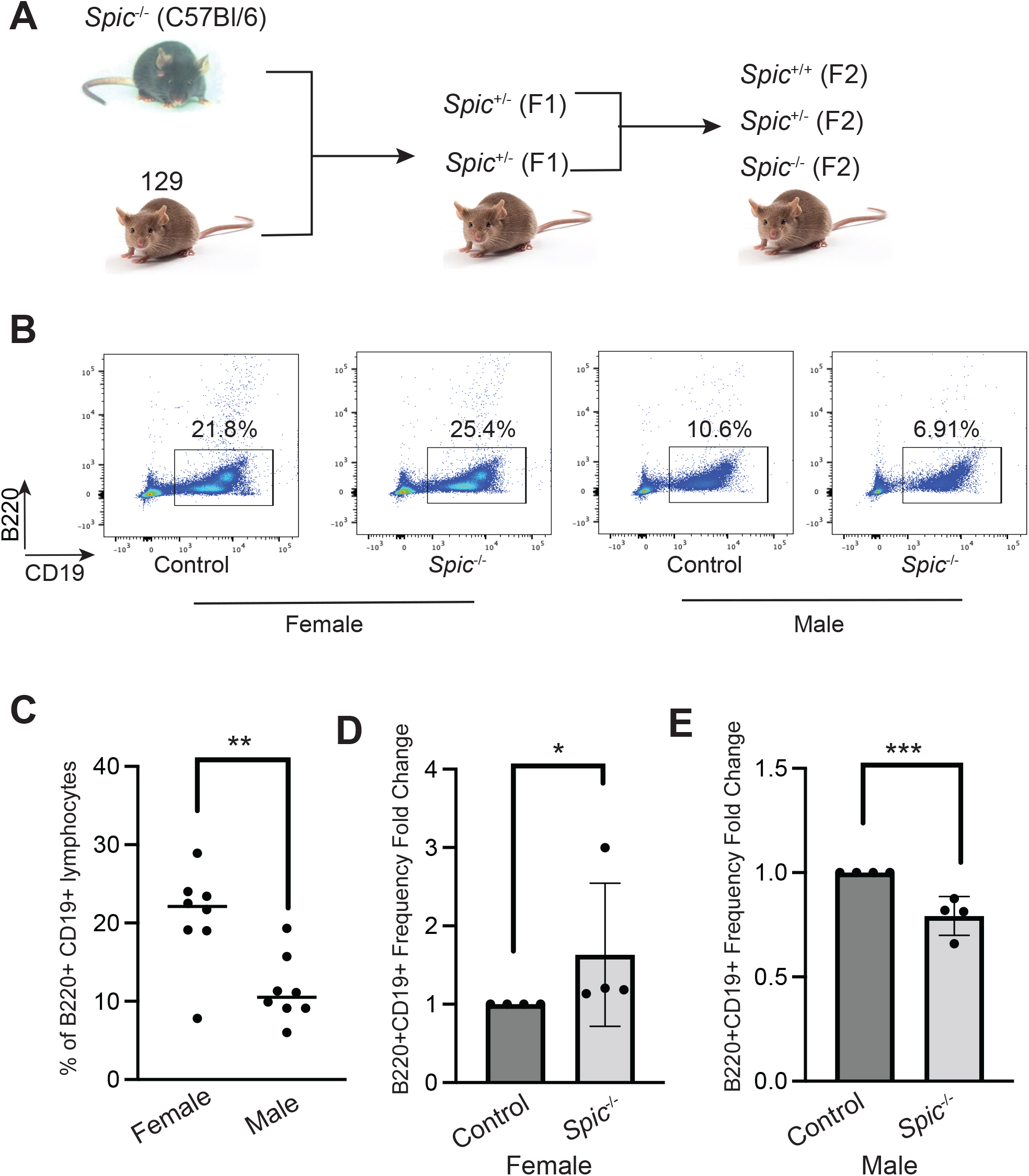
Generation of *Spic*^-/-^ mice. (**A**) Schematic of breeding strategy. (**B**) Representative flow cytometric analysis of B220^+^ CD19^+^ B cell frequencies in the bone marrow of mice of the indicated genotypes. (**C**) Frequencies of B220^+^ CD19^+^ B cells in the bone marrow of female and male *Spic*^+/+^ F2 mice. **p<0.01 using Student’s t-test. (**D**) Relative frequencies of B220^+^ CD19^+^ B cells in the bone marrow of female *Spic*^+/+^ (control) and *Spic*^-/-^ F2 mice. \**p*<0.05 using one-sample t and Wilcoxon test. (**E**) Relative frequencies of B220^+^ CD19^+^ B cells in the bone marrow of male *Spic*^+/+^ (control) and *Spic*^-/-^ F2 mice. \**p*<0.05 using one-sample t and Wilcoxon test.

**Table 1.** Expected and actual frequencies of *Spic*^-/-^ mice at 14.5 days post implantation or 3 weeks after birth from F2 *Spic*^+/-^ intercrosses

| F1 Genetic background | Mouse Age upon Genotyping Characterization | Genotype | No. of Mice Produced | Expected Frequency-*a* (%) | Actual Frequency-*b* (%) |
| --- | --- | --- | --- | --- | --- |
| C57BL/6 | 14.5 d.p.i. | Spic+/+ | 6 | 25 | 15.4 |
|  |  | Spic+/- | 31 | 50 | 79.5 |
|  |  | Spic-/- | 2 | 25 | 5 |
| C57BL/6 | 3 wks | Spic+/+ | 5 | 25 | 21.7 |
|  |  | Spic+/- | 18 | 50 | 78.3 |
|  |  | Spic-/- | 0 | 25 | 0 |
| C57BL and 129 | 3 wks | Spic+/+ | 99 | 25 | 30 |
|  |  | Spic+/- | 182 | 50 | 55.2 |
|  |  | Spic-/- | 49 | 25 | 14.8 |

*a*-based on Mendelian ratio
*b*- based on the caculation from total mice within the group

There are substantial sex differences between frequencies of lymphocytes in various mouse strains (39). We found that female B6/129 F2 mice had increased frequencies of CD19^+^ B220^+^ B cells in the bone marrow compared to male mice (Fig. 3C). Interestingly, *Spic*^-/-^ mice had higher frequencies of B cells in females (Fig. 3D) and lower frequencies of B cells in males (Fig. 3E) compared to littermate control mice. This result suggests a sex-specific effect of Spi-C on B cell development.

The frequency of bone marrow B cell development stages was determined using the Hardy scheme in which B cell development is separated into fractions A-F (40). No significant differences were detected between *Spic*^-/-^ female mice and *Spic^+/+^* or *Spic*^+/-^ female control mice (Fig. 4A, B) or between male mice of the same genotypes (data not shown). Next, *Ig*κ V-J rearrangements were determined. RT-PCR analysis of RNA prepared from enriched fraction D BM pre-B cells showed similar frequencies of Vκ-Jκ rearrangements in *Spic*^-/-^ mice and *Spic*^+/+^ control mice (Fig. 4C). To perform quantitative analysis, Vκ-Jκ5 PCR products were sequenced (Fig. 4D) and used to design qPCR primers. RT-qPCR analysis of Vκ-Jκ5 rearrangements revealed no quantitative differences in fraction D pre-B cells (Fig. 4E) or fraction F immature B cells (Fig. 4F) from *Spic*^-/-^ mice and *Spic^+/+^* control mice. In summary, these results suggest that loss of Spi-C function does not impair early B cell development.

**FIGURE 4.**
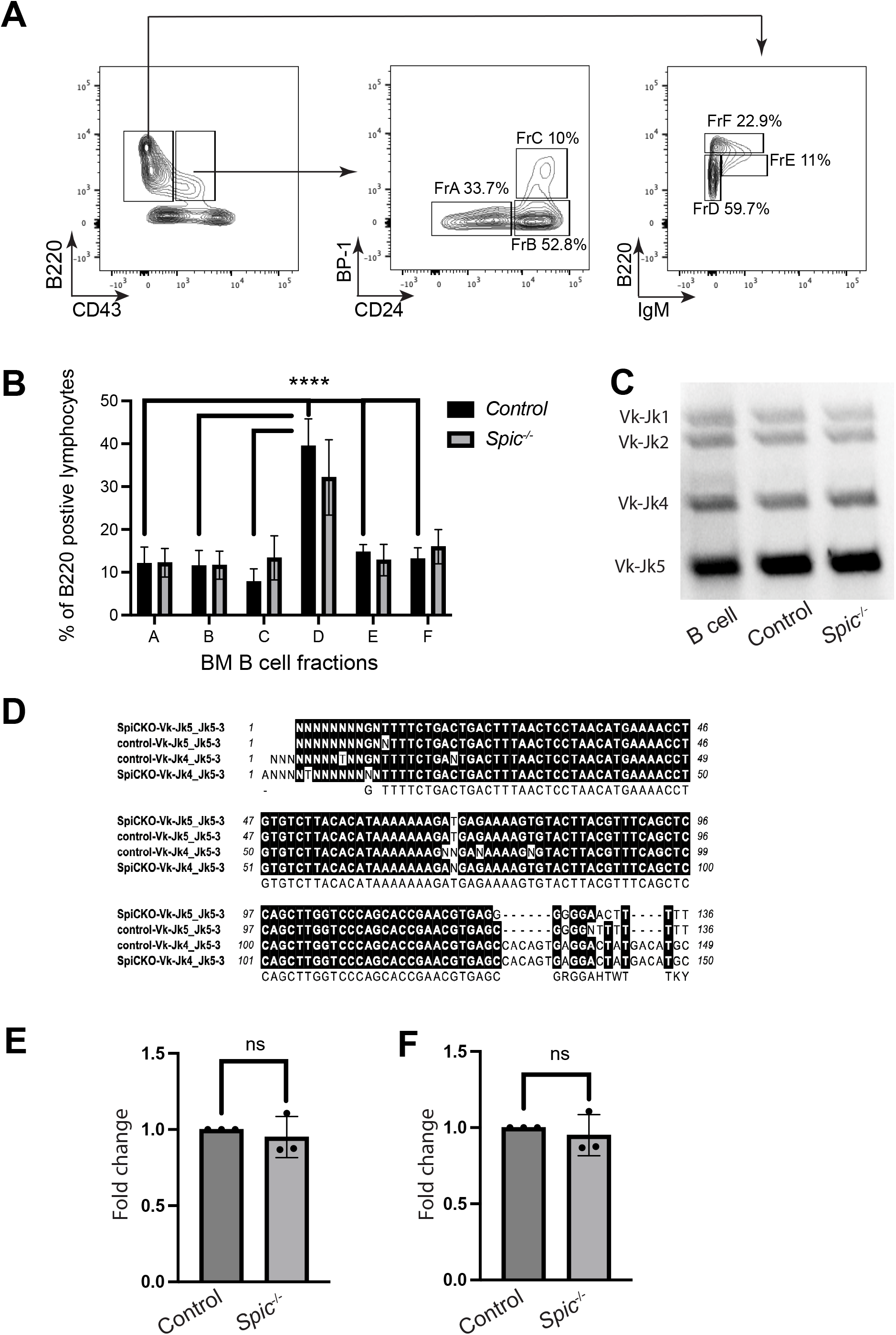
Early B cell development is not altered in *Spic*^-/-^ mice. (**A**) Gating strategy for determination of bone marrow developmental stages. Cell surface markers are indicated on the x and y-axes and boxes are labelled with developmental stage identifications (Fractions A-F) and representative frequencies within each gate. (**B**) Frequencies of bone marrow B cell fractions A-F (x-axis) expressed as per cent of total bone marrow B220^+^ lymphocytes (y-axis). Frequencies of female *Spic*^+/+^ or *Spic*^+/-^ (control) mice and *Spic*^-/-^ F2 mice are indicated in the legend. (**C**) PCR detection of *Ig*κ V-J rearrangements in genomic DNA prepared from B220^+^ CD19^+^ B cells enriched from the bone marrow of female F2 mice of the indicated genotype enriched by cell sorting. (**D**) Representative DNA sequences of amplification products of Vκ-Jκ5 rearrangements from two control and two *Spic*^-/-^ mice. The Vκ forward primer was used as the sequencing primer. (**E**) Expression relative to control of Vκ-Jκ5 rearrangements detected in bone marrow B cells enriched from female *Spic*^+/+^ (control) and *Spic*^-/-^ F2 mice is shown on the y-axis. NS indicates not significant using one-sample t and Wilcoxon test.

### Deletion of the Spic gene in mice leads to elevated levels of Rag1 and Ig kappa sterile transcripts

The low frequency of bone marrow B cells in male B6/129 F2 mice *Spic*^-/-^ mice made enrichment of B cells from male mice by cell sorting technically unfeasible. We therefore enriched total bone marrow CD19^+^ B220^+^ B cells from female B6/129 F2 *Spic*^-/-^ mice, as well as female littermate *Spic*^+/+^ or *Spic*^+/-^ mice as controls. RNA was prepared from enriched B cells for RT-qPCR analysis of gene expression. *Spic* mRNA transcripts were undetectable in *Spic*^-/-^ BM B cells, as expected (Fig. 5A). *Rag1* and *Glk1* were expressed at significantly higher levels in B cells from *Spic*^-/-^ female mice compared to *Spic*^+/+^ or *Spic*^+/-^ female mice (Fig. 5B, D). *Rag2* trended towards higher levels of expression in *Spic*^-/-^ female mice compared to *Spic*^+/+^ or *Spic*^+/-^ female mice (Fig. 5C). These results suggested that Spi-C controls levels of *Rag1* and *Glk1* transcripts in bone marrow B cells.

**FIGURE 5.**
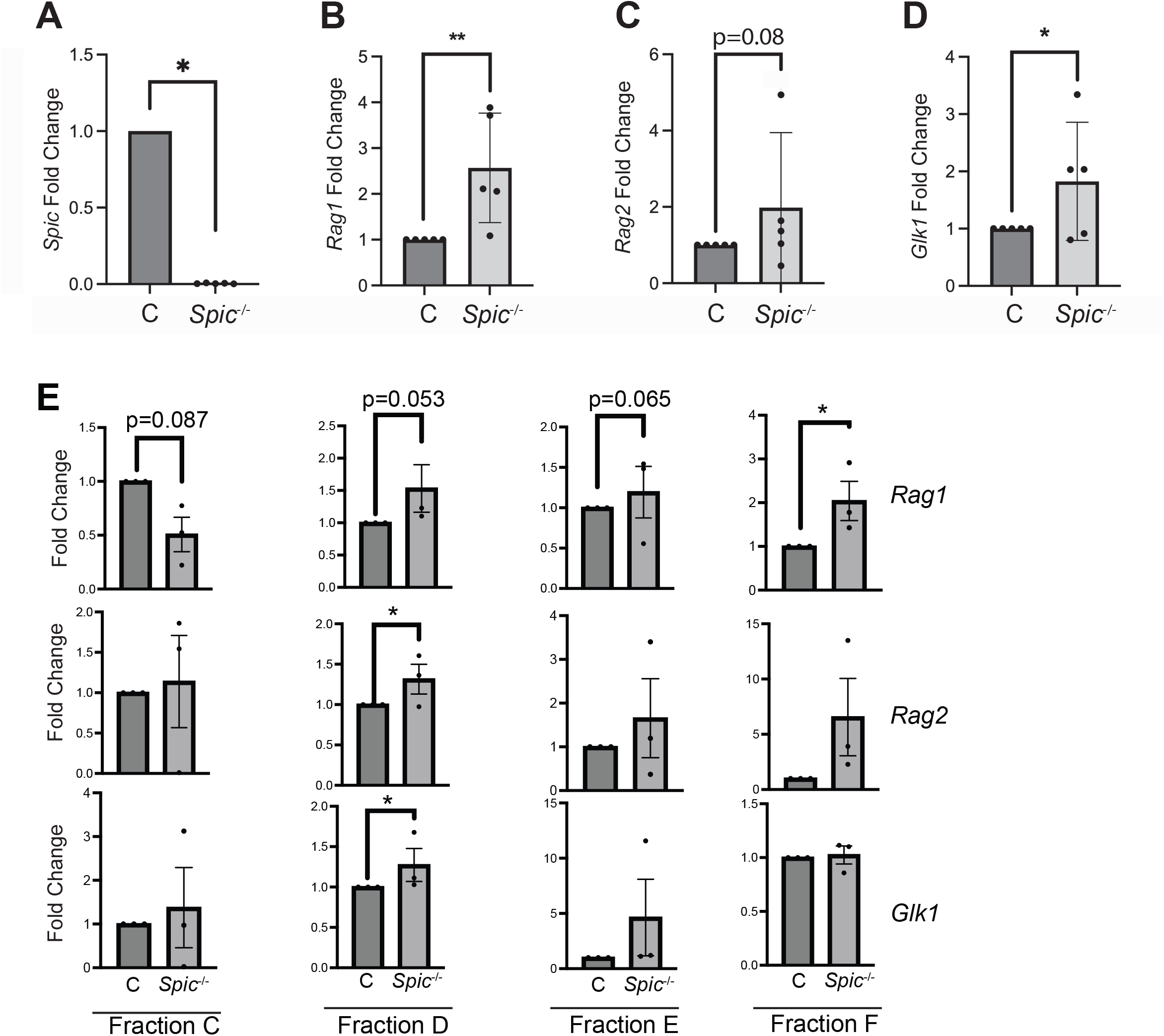
Increased relative levels of *Rag1* and *Glk1* transcripts in bone marrow B cells from *Spic*^-/-^ mice. (**A-D**) RNA was prepared from B220^+^ CD19^+^ B cells enriched from the bone marrow of female F2 *Spic*^-/-^ or control mice. RT-qPCR was performed for the genes indicated on the y-axis. Relative expression (gene/β-*actin* ratios) are indicated on the y-axis. \**p*<0.05, **p<0.01 using one-sample t and Wilcoxon test. (**E**) RNA was prepared from bone marrow fractions C-F enriched from the bone marrow of female F2 *Spic*^-/-^ or control mice. RT-qPCR was performed for the genes indicated on the right side. Expression relative to control is indicated on the y-axis. \**p*<0.05 using one-sample t and Wilcoxon test.

Next, fraction C, D, E, and F cells were enriched by cell sorting from *Spic*^-/-^ female mice and *Spic*^+/+^ or *Spic*^+/-^ female mice and RNA was prepared to determine gene expression using RT-qPCR. *Rag1* was expressed at higher levels in *Spic*^-/-^ fraction D cells and fraction F cells than in controls (Fig. 5E). *Rag2* and *Glk1* were expressed at higher levels in *Spic*^-/-^ fraction D cells than in control fraction D cells. The majority of fraction D cells represent pre-B cells that have stopped proliferating in response to IL-7 and have initiated rearrangement of Igκ, but do not yet express a BCR (40). In agreement with this observation, we found that Fraction D cells expressed higher levels of *Rag* and *Glk1* mRNA transcripts than Fraction C, E, or F cells; and the difference was more pronounced in *Spic*^-/-^ B cells than in WT B cells (Fig. 6). In summary, our results suggest that Spi-C represses *Rag* and *Glk1* transcription starting at the pre-B (Fraction D) cell stage.

**FIGURE 6.**
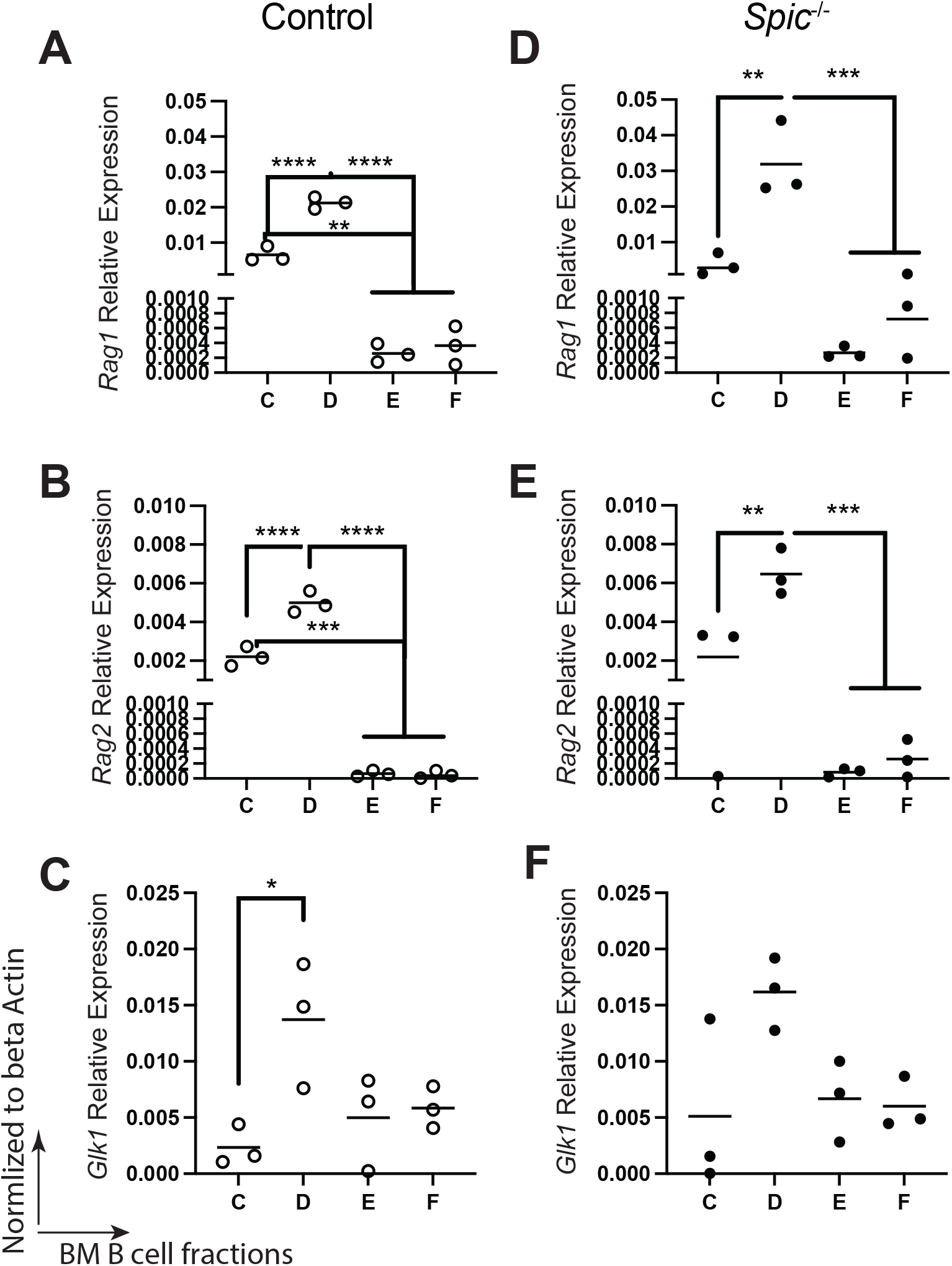
Analysis of *Rag1, Rag2* and *Glk1* mRNA transcript levels in fraction C-F bone marrow B cells from *Spic*^-/-^ mice. For panels **A-F,** RNA was prepared from bone marrow fractions C-F enriched from the bone marrow of female F2 *Spic*^-/-^ or control mice. RT-qPCR was performed for the genes indicated on the y-axis in each panel. Relative expression (gene/β-*actin* ratios) are indicated on the y-axis. \**p*<0.05 using one-way ANOVA.

### PU.1 activates Ig kappa germline and Rag1 transcription

PU.1 and Spi-B are highly related to Spi-C in the ETS transcription factor family. We previously showed that PU.1 is an activator of *Ig*κ transcription and rearrangement during B cell development (24). To further explore this observation, we made use of the i660BM pre-B cell line in which PU.1 is doxycycline-inducible (33). Doxycycline induction of PU.1 in i660BM cells for 48 hr (Fig. 7A) increased *Rag1* and *Rag2* mRNA transcript levels (Fig 7B, C). This result suggests that PU.1 is a direct activator of *Rag* transcription. Next, we looked at mice in which PU.1 is inducibly deleted under control of the *Mb1* (*Cd79a*) gene starting at the pro-B stage (Mb1-CreΔPB mice) (24). These mice are also germline null for the *Spib* gene. We found that *Rag1* and *Glk1* mRNA transcripts were decreased in Fraction D cells enriched from Mb1-CreΔPB mice compared to *Spib*^-/-^ control mice (Fig. 7D-G). These data suggest that PU.1 functions as an activator of *Ig*κ and *Rag1* transcription and *Ig*κ rearrangement. Taken together with the previous results, these data suggest that Spi-C and PU.1 act in an opposite manner to regulate *Ig*κ and *Rag1* transcription and *Ig*κ rearrangement starting at the small pre-B cell stage.

**FIGURE 7.**
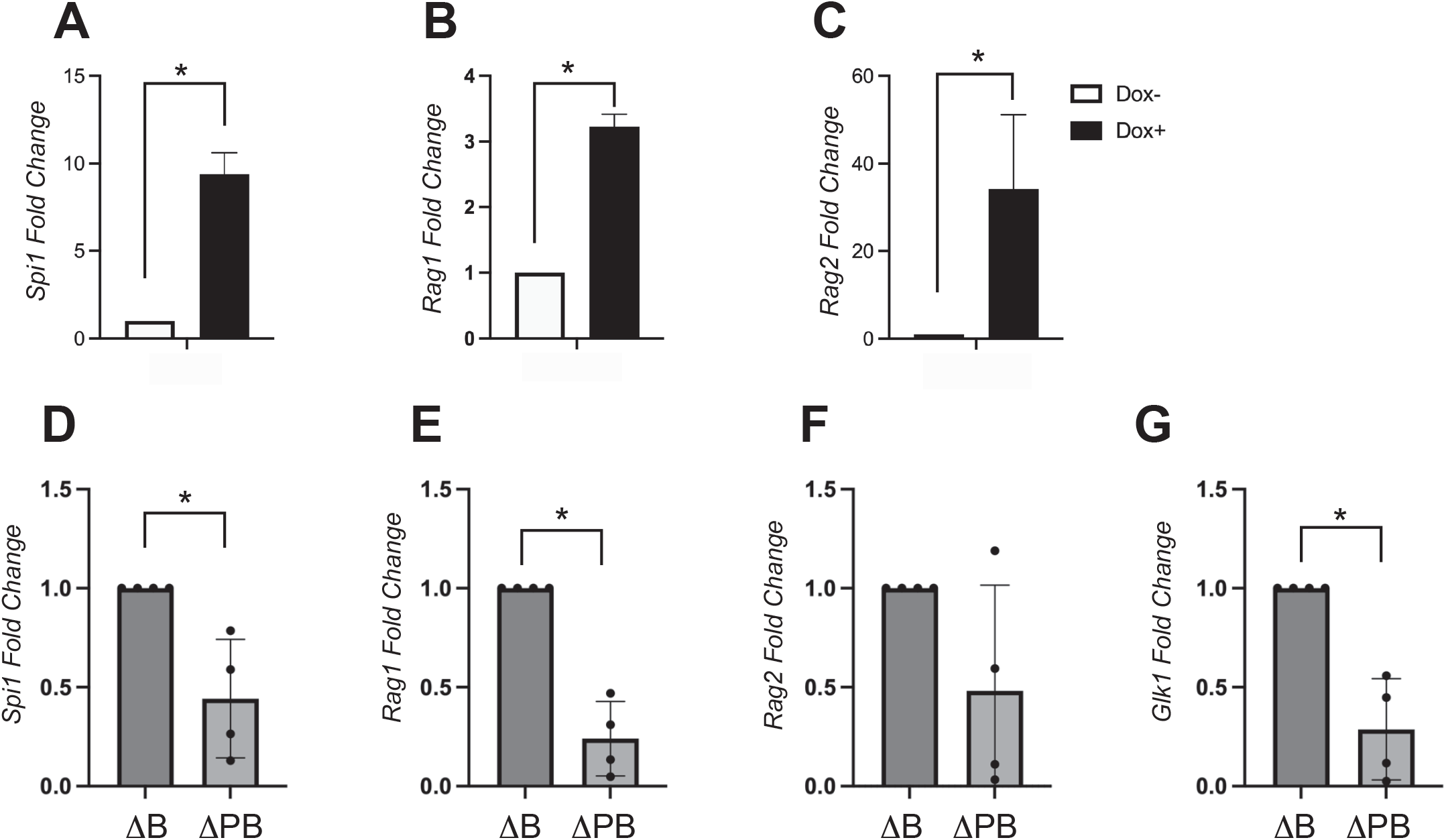
PU.1 activates *Ig* kappa germline and *Rag1* transcription. (**A-C**) PU.1-inducible i660BM cells were treated with 70 ng/ml doxycycline (Dox+) or without (Dox-) for 48 hr followed by preparation of RNA. RT-qPCR was performed to determine relative expression levels of the genes indicated on the y-axis normalized to β-actin levels. \**p*<0.05 using one sample t and Wilcoxon test. (**D-G**) PU.1 is required for gene expression in fraction D pre-B cells. Fraction D pre-B cells were enriched by cell sorting from bone marrow of female Mb1-CreΔPB mice lacking both PU.1 and Spi-B (ΔPB) or *SpiN*^-/-^ mice as control (ΔB). RT-qPCR was performed to determine relative levels of the genes indicated on the y-axis. \**p*<0.05 using one sample t and Wilcoxon test.

### Direct interaction of PU.1 and Spi-C with a site in the Rag locus

To determine if Spi-C regulates *Rag* transcription directly by interaction with a binding site in a regulatory region, we performed anti-Spi-C chromatin immunoprecipitation (ChIP). 3XFLAG-Spi-C was induced by doxycycline in i38B9 cells for 48 hr, after which cells were formaldehyde fixed and chromatin prepared for immunoprecipitation. ChIP was performed using anti-FLAG antibody as described in Materials & Methods. First, qPCR analysis was performed on a previously identified Spi-C binding site (23) in the first intron of the *Bach2* gene (Fig. 8A). Using qPCR analysis, Spi-C binding at *Bach2* Region of Interest 1 (ROI1) was increased 40-fold compared to binding at a negative control region (NCR) (Fig. 8B). Next, qPCR analysis was performed to determine Spi-C binding at a previously identified PU.1 binding site within the *Rag* locus (24) (Fig. 8C). Four independent biological replicate experiments found 8-45 fold increased binding of Spi-C at the *Rag* peak compared to a negative control region (NCR) in the same locus (Fig. 8D-G). In summary, Spi-C and PU.1 interact with a binding site located directly upstream of the *Rag1* gene, suggesting that Spi-C and PU.1 regulate Rag transcription directly.

**FIGURE 8.**
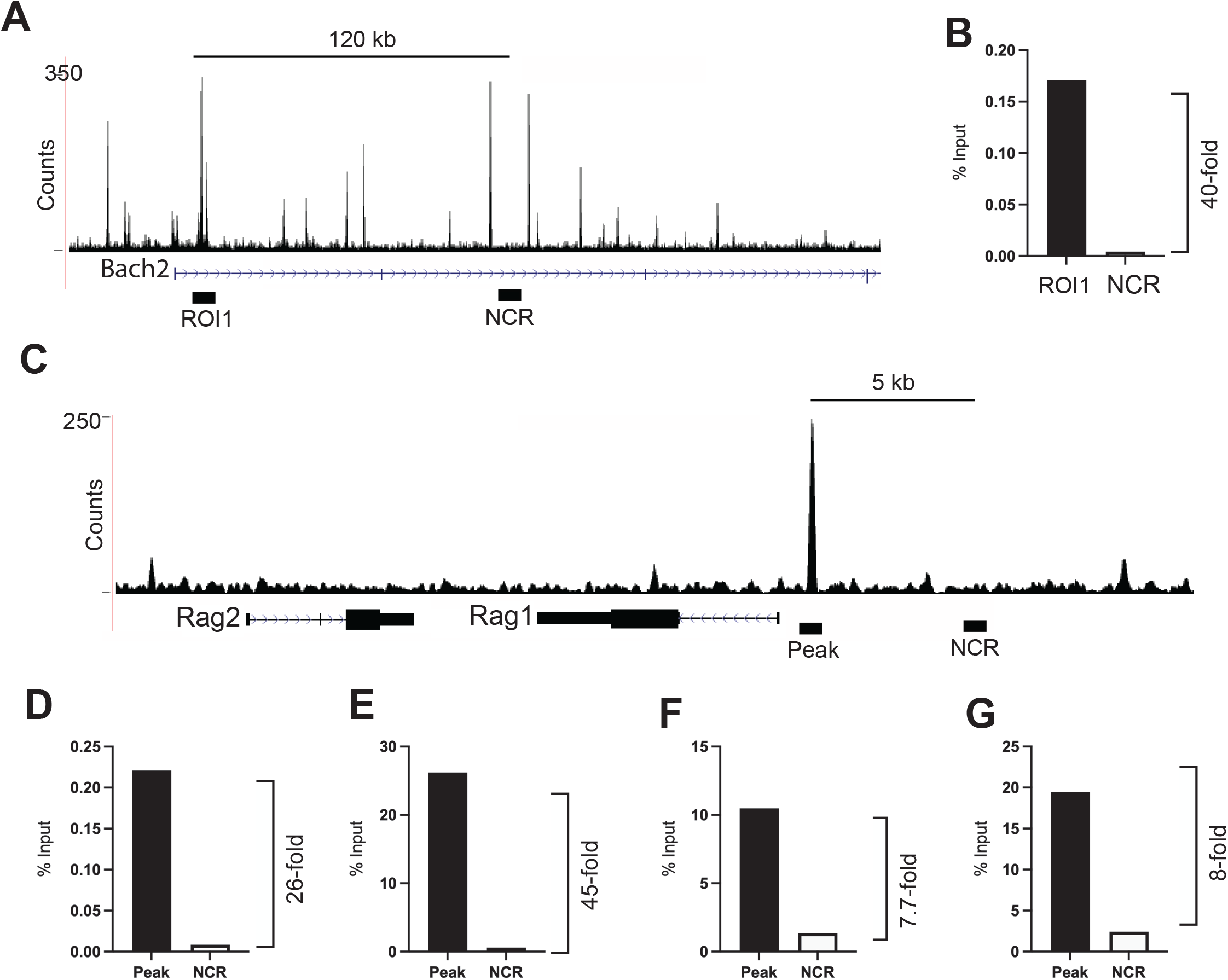
Direct interaction of PU.1 and Spi-C with a site in the *Rag* locus. (**A**) UCSC genome browser track of PU.1 ChIP-seq data showing the *Bach2* gene. Black bars indicate region of interest 1 (ROI1) and negative control region (NCR). **(B**) Enrichment of Spi-C at ROI1 relative to the NCR. Anti-Spi-C ChIP was performed in i38B9 cells as described in Materials & Methods and is shown as per cent of input (y-axis). (**C**) USCS genome browser track of PU.1 ChIP-seq data showing the *Rag* locus. Black bars indicate a region of PU.1 binding (Peak) and a negative control region (NCR). (**D**-**G**) Enrichment of Spi-C at the PU.1 binding site (Peak) relative to NCR. D-G represent four biological replicates. Fold enrichment is shown as per cent of input).

## Discussion

In this study, we investigated the ability of the ETS transcription factor Spi-C to negatively regulate *Ig*κ light chain recombination. Using an inducible expression system in a pre-B cell line, we found that Spi-C negatively regulated *Ig*κ rearrangement, *Ig*κ transcript levels, and *Rag1* transcript levels. By analyzing bone marrow-derived B cells from *Spic*^-/-^ mice, we found that *Ig*κ and *Rag1* transcript levels were negatively regulated by Spi-C in small pre-B cells. In contrast, we found that *Ig*κ and *Rag1* transcript levels were activated by PU.1 in small pre-B cells. Using chromatin immunoprecipitation analysis, we identified a binding site for PU.1 and Spi-C located 5’ of the *Rag1* promoter. We conclude that Spi-C and PU.1 are counterregulators of *Ig*κ light chain recombination and *Rag1* transcription in small pre-B cells.

Double-stranded DNA breaks (DSBs) induced at the *Ig*κ light chain locus in small pre-B cells, have been demonstrated to negatively feed back on *Rag* and *Ig*κ transcription to promote allelic exclusion at the *Ig*κ locus (16). The signaling pathway induced by DSBs to promote allelic exclusion requires ataxia telangiectasia mutated (ATM) kinase and the NF-κB essential modulator (Nemo) protein (16). This signaling pathway activates several genes including *Spic* (18). Spi-C is highly related to PU.1 and Spi-B and can interact with and/or compete for similar binding sites in the genome of B cells and macrophages (29, 30, 41). Spi-C can repress *Ig*κ recombination and cell division by interacting with BCLAF1 to displace PU.1, resulting in repression of the *Syk*, *Blnk*, and *Ig*κ genes (18, 19) Although *Rag1* transcription was shown to be negatively regulated by DSBs, it has not been clear what factors mediate this transcriptional repression (42). Our study now demonstrates that Spi-C is a direct negative regulator of *Rag1* transcription in pre-B cells.

PU.1 has been shown in multiple studies to function primarily as an activator of transcription (43). In addition, PU.1 functions as a pioneer protein, having the ability to interact with binding sites in closed chromatin and recruit histone acetyltransferases to open chromatin sites (44). In our previous studies and in the study by Soodgupta et al (19), Spi-C appeared to function as a negative regulator of gene transcription primarily by displacing PU.1 from binding sites. Chromatin immunoprecipitation studies suggest that most Spi-C binding sites are also PU.1 binding sites, whereas there are fewer unique Spi-C binding sites (29). It remains to be determined whether Spi-C interaction with target sites requires prior PU.1 binding or pioneer activity.

The *Ig*κ 3’ enhancer contains an important ETS transcription factor binding site that regulates B cell versus T cell specificity of Vk-Jk joining (7). PU.1 was shown to interact with this site to promote *Ig*κ transcription and accessibility (45, 46). Because of functional redundancy and complementarity with PU.1, Spi-B very likely also regulates *Ig*κ transcription through this site (24). Spi-C was shown to displace PU.1 interaction with binding sites in the *Ig*κ locus (19). Therefore, Spi-C likely negatively regulates the *Glk1* mRNA transcript by direct interaction with PU.1 interaction sites in the *Ig*κ locus.

In all experiments performed in this study, *Rag1* was affected to a greater degree than *Rag2* by Spi-C ectopic expression or deletion. We speculate that this is because of the proximity of the *Rag1* transcription start site to the binding site that we identified for PU.1/Spi-C. Mechanisms of transcriptional regulation of the *Rag* locus have not been well-studied, with only the identification of the ERag enhancer reported for many years (14). However, a more detailed study of *Rag* transcriptional regulation was reported in 2020 by Miyazaki *et al*, using ATAC-seq to identify additional regulatory regions named R-TEn, R2B, R1pro, and R1B (15). R1pro represents the *Rag1* promoter region and was shown to interact with multiple transcription factors including E2A, Pax5, ETS1, Ikaros, and IRF4. We have now shown that Spi-C and PU.1 also interact with the R1pro region, increasing the probability that this is an important regulatory region for the *Rag1* gene.

We speculate that in small B cells, PU.1 activates *Ig*κ and *Rag* transcription using its pioneer activity. PU.1 is a strong transcriptional activator of *Ig*κ (24) and as we show here, *Rag1*, to promote *Ig*κ recombination. Following Rag1/2-mediated DNA cleavage at one *Ig*κ allele, The *Spic* gene is activated, resulting in Spi-C interaction with BCLAF1 to displace PU.1 from target genes in small pre-B cells. Spi-C expression may persist into the transitional-1 stage to modulate BCR signaling. In this manner, PU.1 and Spi-C are proposed to function as counterregulators of *Ig*κ allelic exclusion; and are potentially involved in regulation of BCR signaling and receptor editing. In summary, our experiments support the idea that Spi-C and PU.1 counterregulate *Ig*κ transcription and *Rag1* transcription to effect *Ig*κ recombination in small pre-B cells.

## Acknowledgements

We thank Dr. Malay Haldar (Perelman School of Medicine, University of Pennsylvania) for advice regarding breeding and genotyping of *Spic*^-/-^ mice. We thank Kristin Chadwick and the London Regional flow cytometry facility for assistance with cell sorting.

## Figure Legends

**Supplemental Table 1.**
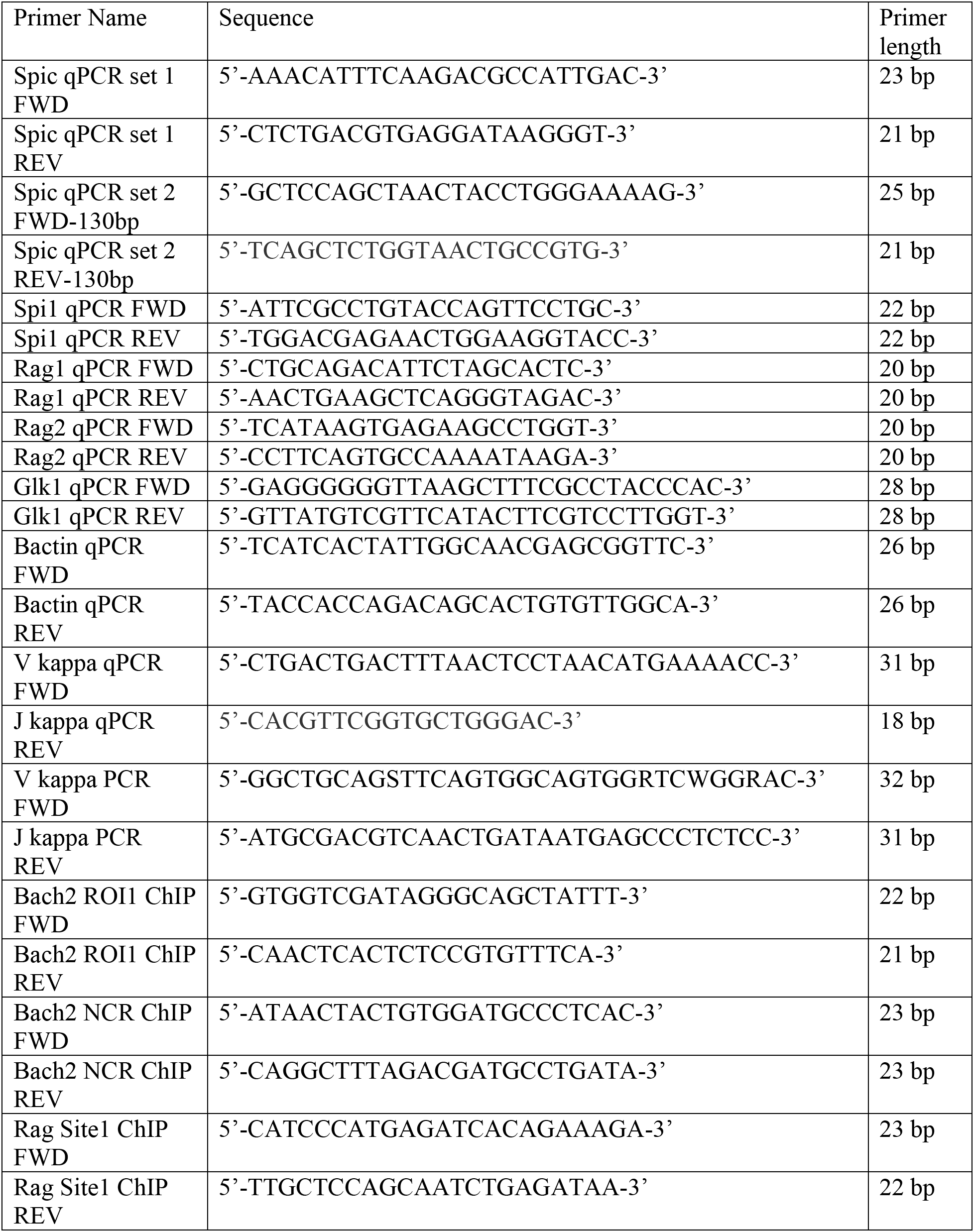
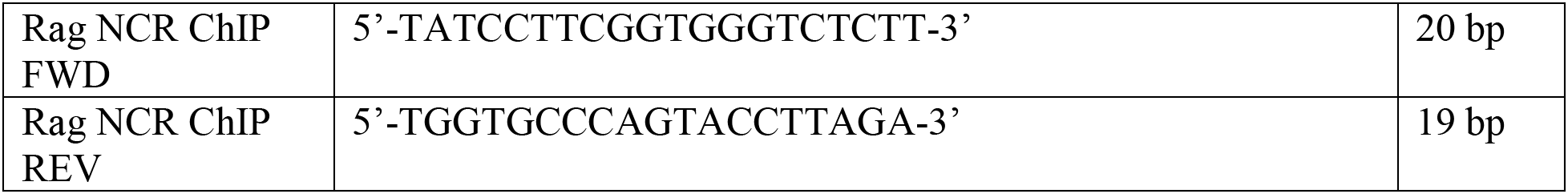
Primer Sequences.

**SUPPLEMENTAL FIGURE 1.**
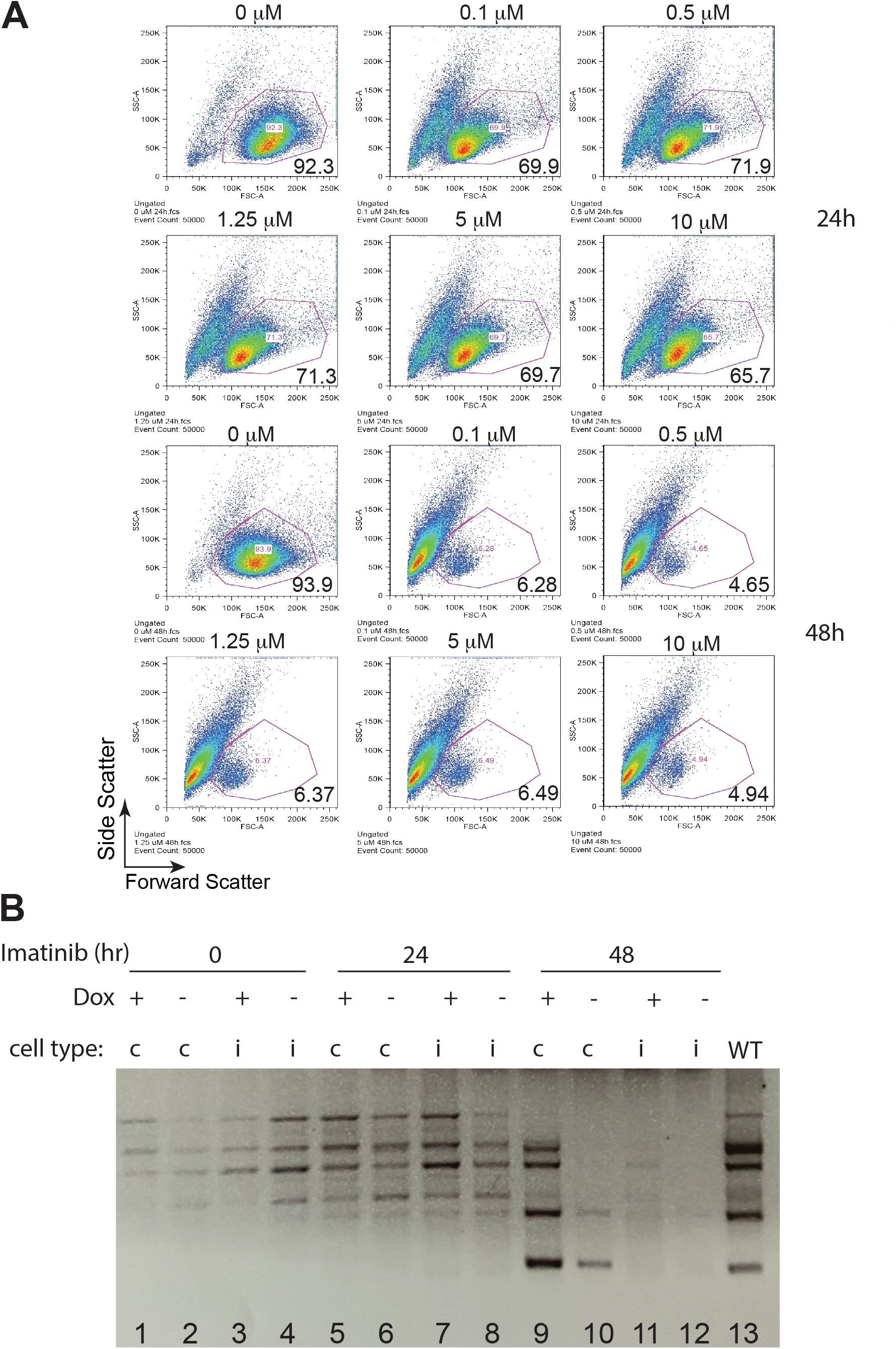
Characterization of Spi-C inducible i38B9 cells. (**A**) Reduction in cell size following imatinib treatment. Inducible 38B9 cells (i38B9 cells) were treated with various concentrations of imatinib (shown on top of dot plots) for 24 hr or 48 hr (indicated on right) followed by flow cytometric analysis of light scatter. Forward scatter is on x-axis, side scatter on y-axis. The frequency of high forward scatter cells is indicated inside the plot. (**B**) Inhibition of *Ig*κ V-J rearrangement 24 or 48 hr after imatinib treatment and Spi-C induction. PCR analysis was performed on genomic DNA prepared from cells treated with 0.5 μM imatinib at times indicated on the top as well as 1.5 μg/ml doxycycline. “A” cell type is 38B9 cells; “B” cell type is i38B9. Positive control (WT) is from genomic DNA prepared from wild-type splenic B cells. \**p*<0.05 and ** *p*<0.01 using two-way ANOVA.

## References

1. Clark, M. R., M. Mandal, K. Ochiai, and H. Singh. 2014. Orchestrating B cell lymphopoiesis through interplay of IL-7 receptor and pre-B cell receptor signalling. Nat Rev Immunol 14: 69–80.

2. Hamel, K. M., M. Mandal, S. Karki, and M. R. Clark. 2014. Balancing Proliferation with Igk Recombination during B-lymphopoiesis. Front Immunol 5.

3. Karki, S., S. Banerjee, K. Mclean, A. Dinner, and M. R. Clark. 2019. Transcription factories in Igκ allelic choice and diversity. Adv Immunol 141: 33–49.

4. Inlay, M., F. W. Alt, D. Baltimore, and Y. Xu. 2002. Essential roles of the kappa light chain intronic enhancer and 3’ enhancer in kappa rearrangement and demethylation. Nat Immunol 3: 463–468.

5. van Ness, B. G., M. Weigert, C. Coleclough, E. L. Mather, D. E. Kelley, and R. P. Perry. 1981. Transcription of the unrearranged mouse C kappa locus: sequence of the initiation region and comparison of activity with a rearranged V kappa-C kappa gene. Cell 27: 593–602.

6. Schlissel, M. S., and D. Baltimore. 1989. Activation of immunoglobulin kappa gene rearrangement correlates with induction of germline kappa gene transcription. Cell 58: 1001–7.

7. Hiramatsu, R., K. Akagi, M. Matsuoka, K. Sakumi, H. Nakamura, L. Kingsbury, C. David, R. R. Hardy, K. Yamamura, and H. Sakano. 1995. The 3’ enhancer region determines the B/T specificity and pro-B/pre-B specificity of immunoglobulin V kappa-J kappa joining. Cell 83: 1113–1123.

8. Ochiai, K., M. Maienschein-Cline, M. Mandal, J. R. Triggs, E. Bertolino, R. Sciammas, A. R. Dinner, M. R. Clark, and H. Singh. 2012. A self-reinforcing regulatory network triggered by limiting IL-7 activates pre-BCR signaling and differentiation. Nat Immunol 13: 300–307.

9. Mandal, M., S. E. Powers, M. Maienschein-Cline, E. T. Bartom, K. M. Hamel, B. L. Kee, A. R. Dinner, and M. R. Clark. 2011. Epigenetic repression of the Igk locus by STAT5-mediated recruitment of the histone methyltransferase Ezh2. Nat Immunol 12: 1212–1220.

10. Teng, G., and D. G. Schatz. 2015. Regulation and Evolution of the RAG Recombinase. Adv Immunol 128: 1–39.

11. Papaemmanuil, E., I. Rapado, Y. Li, N. E. Potter, D. C. Wedge, J. Tubio, L. B. Alexandrov, P. van Loo, S. L. Cooke, J. Marshall, I. Martincorena, J. Hinton, G. Gundem, F. W. van Delft, S. Nik-Zainal, D. R. Jones, M. Ramakrishna, I. Titley, L. Stebbings, C. Leroy, A. Menzies, J. Gamble, B. Robinson, L. Mudie, K. Raine, S. O’Meara, J. W. Teague, A. P. Butler, G. Cazzaniga, A. Biondi, J. Zuna, H. Kempski, M. Muschen, A. M. Ford, M. R. Stratton, M. Greaves, and P. J. Campbell. 2014. RAG-mediated recombination is the predominant driver of oncogenic rearrangement in ETV6-RUNX1 acute lymphoblastic leukemia. Nat Genet 46: 116–125.

12. Lin, W. C., and S. Desiderio. 1994. Cell cycle regulation of V(D)J recombination-activating protein RAG-2. Proc Natl Acad Sci U S A 91: 2733–2737.

13. Kuo, T. C., and M. S. Schlissel. 2009. Mechanisms controlling expression of the RAG locus during lymphocyte development. Curr Opin Immunol 21: 173–8.

14. Hsu, L.-Y., J. Lauring, H.-E. Liang, S. Greenbaum, D. Cado, Y. Zhuang, and M. S. Schlissel. 2003. A conserved transcriptional enhancer regulates RAG gene expression in developing B cells. Immunity 19: 105–17.

15. Miyazaki, K., H. Watanabe, G. Yoshikawa, K. Chen, R. Hidaka, Y. Aitani, K. Osawa, R. Takeda, Y. Ochi, S. Tani-ichi, T. Uehata, O. Takeuchi, K. Ikuta, S. Ogawa, G. Kondoh, Y. C. Lin, H. Ogata, and M. Miyazaki. 2020. The transcription factor E2A activates multiple enhancers that drive Rag expression in developing T and B cells. Sci Immunol 5: eabb1455.

16. Fisher, M. R., A. Rivera-Reyes, N. B. Bloch, D. G. Schatz, and C. H. Bassing. 2017. Immature Lymphocytes Inhibit Rag1 and Rag2 Transcription and V(D)J Recombination in Response to DNA Double-Strand Breaks. J Immunol 198: 2943–2956.

17. Raczkowski, H. L., and R. P. DeKoter. 2021. Lineage-instructive functions of the E26 - transformation-specific-family transcription factor Spi-C in immune cell development and disease. WIREs Mechanisms of Disease 13: e1519.

18. Bednarski, J. J., R. Pandey, E. Schulte, L. S. White, B. R. Chen, G. J. Sandoval, M. Kohyama, M. Haldar, A. Nickless, A. Trott, G. Cheng, K. M. Murphy, C. H. Bassing, J. E. Payton, and B. P. Sleckman. 2016. RAG-mediated DNA double-strand breaks activate a cell type-specific checkpoint to inhibit pre-B cell receptor signals. J Exp Med 213: 209–223.

19. Soodgupta, D., L. S. White, W. Yang, R. Johnston, J. M. Andrews, M. Kohyama, K. M. Murphy, N. Mosammaparast, J. E. Payton, and J. J. Bednarski. 2019. RAG-Mediated DNA Breaks Attenuate PU.1 Activity in Early B Cells through Activation of a SPIC-BCLAF1 Complex. Cell Rep 29: 829–843.e5.

20. Kohyama, M., W. Ise, B. T. Edelson, P. R. Wilker, K. Hildner, C. Mejia, W. A. Frazier, T. L. Murphy, and K. M. Murphy. 2009. Role for Spi-C in the development of red pulp macrophages and splenic iron homeostasis. Nature 457: 318–321.

21. Haldar, M., M. Kohyama, A. Y. L. So, W. Kc, X. Wu, C. G. Briseño, A. T. Satpathy, N. M. Kretzer, H. Arase, N. S. Rajasekaran, L. Wang, T. Egawa, K. Igarashi, D. Baltimore, T. L. Murphy, and K. M. Murphy. 2014. Heme-mediated SPI-C induction promotes monocyte differentiation into iron-recycling macrophages. Cell 156: 1223–1234.

22. Raczkowski, H. L., L. S. Xu, W. C. Wang, and R. P. DeKoter. 2022. The E26 Transformation–Specific Family Transcription Factor Spi-C Is Dynamically Regulated by External Signals in B Cells. Immunohorizons 6: 104.

23. Laramée, A. S., H. Raczkowski, P. Shao, C. Batista, D. Shukla, L. Xu, S. M. M. Haeryfar, Y. Tesfagiorgis, S. Kerfoot, and R. DeKoter. 2020. Opposing Roles for the Related ETS-Family Transcription Factors Spi-B and Spi-C in Regulating B Cell Differentiation and Function. Front Immunol 11.

24. Batista, C. R., S. K. H. Li, L. S. Xu, L. A. Solomon, and R. P. DeKoter. 2017. PU.1 Regulates Ig Light Chain Transcription and Rearrangement in Pre-B Cells during B Cell Development. J Immunol 198: 1565–1574.

25. Sokalski, K. M., S. K. Li, I. Welch, H. A. Cadieux-Pitre, M. R. Gruca, and R. P. DeKoter. 2011. Deletion of genes encoding PU.1 and Spi-B in B cells impairs differentiation and induces pre-B cell acute lymphoblastic leukemia. Blood 118: 2801–2808.

26. Batista, C. R., M. Lim, A.-S. Laramee, F. Abu-Sardanah, L. S. Xu, R. Hossain, G. I. Bell, D. A. Hess, and R. P. DeKoter. 2018. Driver mutations in Janus kinases in a mouse model of B-cell leukemia induced by deletion of PU.1 and Spi-B. Blood Adv 2: 2798–2810.

27. Zhu, X., B. L. Schweitzer, E. J. Romer, C. E. W. Sulentic, and R. P. DeKoter. 2008. Transgenic expression of Spi-C impairs B-cell development and function by affecting genes associated with BCR signaling. Eur J Immunol 38.

28. Schweitzer, B. L., K. J. Huang, M. B. Kamath, A. V. Emelyanov, B. K. Birshtein, and R. P. DeKoter. 2006. Spi-C has opposing effects to PU.1 on gene expression in progenitor B cells. Journal of Immunology 177.

29. Laramée, A.-S., H. Raczkowski, P. Shao, C. Batista, D. Shukla, L. Xu, S. M. M. Haeryfar, Y. Tesfagiorgis, S. Kerfoot, and R. DeKoter. 2020. Opposing Roles for the Related ETS-Family Transcription Factors Spi-B and Spi-C in Regulating B Cell Differentiation and Function. Front Immunol 11.

30. Li, S. K., L. A. Solomon, P. C. Fulkerson, and R. P. DeKoter. 2015. Identification of a negative regulatory role for Spi-C in the murine B cell lineage. The Journal of Immunology 194: 3978–3807.

31. Alt, F. W., N. Rosenberg, V. Enea, E. Siden, and D. Baltimore. 1982. Multiple immunoglobulin heavy-chain gene transcripts in Abelson murine leukemia virus-transformed lymphoid cell lines. Mol Cell Biol 2: 386–400.

32. Muljo, S. A., and M. S. Schlissel. 2003. A small molecule Abl kinase inhibitor induces differentiation of Abelson virus-transformed pre-B cell lines. Nat Immunol 4: 31–37.

33. Christie, D. A., L. S. Xu, S. A. Turkistany, L. A. Solomon, S. K. Li, E. Yim, I. Welch, G. I. Bell, D. A. Hess, and R. P. DeKoter. 2015. PU.1 Opposes IL-7-Dependent Proliferation of Developing B Cells with Involvement of the Direct Target Gene Bruton Tyrosine Kinase. J Immunol 194: 595–605.

34. Winkler, T. H., F. Melchers, and A. G. Rolink. 1995. Interleukin-3 and interleukin-7 are alternative growth factors for the same B-cell precursors in the mouse. Blood 85: 2045–2051.

35. Xu, L. S., K. M. Sokalski, K. Hotke, D. a Christie, O. Zarnett, J. Piskorz, G. Thillainadesan, J. Torchia, and R. P. DeKoter. 2012. Regulation of B cell linker protein transcription by PU.1 and Spi-B in murine B cell acute lymphoblastic leukemia. J Immunol 189: 3347–54.

36. Morita, S., T. Kojima, and T. Kitamura. 2000. Plat-E: an efficient and stable system for transient packaging of retroviruses. Gene Ther 7: 1063–1066.

37. Cobaleda, C., W. Jochum, and M. Busslinger. 2007. Conversion of mature B cells into T cells by dedifferentiation to uncommitted progenitors. Nature 449: 473–477.

38. Alam, Z., S. Devalaraja, M. Li, T. K. J. To, I. W. Folkert, E. Mitchell-Velasquez, M. T. Dang, P. Young, C. J. Wilbur, M. A. Silverman, X. Li, Y. H. Chen, P. T. Hernandez, A. Bhattacharyya, M. Bhattacharya, M. H. Levine, and M. Haldar. 2020. Counter Regulation of Spic by NF-κB and STAT Signaling Controls Inflammation and Iron Metabolism in Macrophages. Cell Rep 31.

39. Hensel, J. A., V. Khattar, R. Ashton, and S. Ponnazhagan. 2019. Characterization of immune cell subtypes in three commonly used mouse strains reveals gender and strain-specific variations. Laboratory Investigation 99: 93–106.

40. Hardy, R. R., C. E. Carmack, S. A. Shinton, J. D. Kemp, and K. Hayakawa. 1991. Resolution and characterization of pro-B and pre-pro-B cell stages in normal mouse bone marrow. J Exp Med 173: 1213–1225.

41. DeKoter, R. P., M. Geadah, S. Khoosal, L. S. Xu, G. Thillainadesan, J. Torchia, S. S. Chin, and L. A. Garrett-Sinha. 2010. Regulation of follicular B cell differentiation by the related E26 transformation-specific transcription factors PU.1, Spi-B, and Spi-C. J Immunol 185: 7374–7384.

42. Glynn, R. A., and C. H. Bassing. 2022. Nemo-Dependent, ATM-Mediated Signals from RAG DNA Breaks at Igk Feedback Inhibit V κ Recombination to Enforce Igκ Allelic Exclusion. J Immunol 208: 371–383.

43. Turkistany, S. A., and R. P. DeKoter. 2011. The transcription factor PU.1 is a critical regulator of cellular communication in the immune system. Arch Immunol Ther Exp (Warsz) 59: 431–440.

44. Minderjahn, J., A. Schmidt, A. Fuchs, R. Schill, J. Raithel, M. Babina, C. Schmidl, C. Gebhard, S. Schmidhofer, K. Mendes, A. Ratermann, D. Glatz, M. Nützel, M. Edinger, P. Hoffmann, R. Spang, G. Längst, A. Imhof, and M. Rehli. 2020. Mechanisms governing the pioneering and redistribution capabilities of the non-classical pioneer PU.1. Nat Commun 11: 402.

45. Pongubala, J. M., S. Nagulapalli, M. J. Klemsz, S. R. McKercher, R. A. Maki, and M. L. Atchison. 1992. PU.1 recruits a second nuclear factor to a site important for immunoglobulin kappa 3’ enhancer activity. Mol Cell Biol 12: 368–378.

46. Hodawadekar, S., K. Park, M. A. Farrar, and M. L. Atchison. 2012. A developmentally controlled competitive STAT5-PU.1 DNA binding mechanism regulates activity of the Ig kappa E3’ enhancer. J Immunol 188: 2276–2284.

